# CLN5 and CLN3 function as a complex to regulate endolysosome function

**DOI:** 10.1101/2020.12.24.423824

**Authors:** Seda Yasa, Etienne Sauvageau, Graziana Modica, Stephane Lefrancois

## Abstract

CLN5 is a soluble endolysosomal protein whose function is poorly understood. Mutations in this protein cause a rare neurodegenerative disease, Neuronal Ceroid Lipofuscinosis. We previously found that depletion of CLN5 leads to dysfunctional retromer, resulting in the degradation of the lysosomal sorting receptor, sortilin. However, how a soluble lysosomal protein can modulate the function of a cytosolic protein, retromer, is not known. In this work, we show that deletion of CLN5 not only results in retromer dysfunction, but also in impaired endolysosome fusion events. This results in delayed degradation of endocytic proteins and in defective autophagy. CLN5 modulates these various pathways by regulating downstream interactions between CLN3, an endolysosomal integral membrane protein whose mutations also result in Neuronal Ceroid Lipofuscinosis, Rab7A, and a subset of Rab7A effectors. Our data supports a model where CLN3 and CLN5 function as an endolysosome complex regulating various functions.

**Summary Statement:** We have previously demonstrated that CLN3 is required for efficient endosome-to-trans Golgi Network (TGN) trafficking of sortilin by regulating retromer function. In this work, we show that CLN5, which interacts with CLN3, regulates retromer function by modulating key interactions between CLN3, Rab7A, retromer, and sortilin. Therefore, CLN3 and CLN5 serve as endosomal switch regulating the itinerary of the lysosomal sorting receptors.

## Introduction

The Neuronal Ceroid Lipofuscinoses (NCLs) are a group of inherited lysosomal diseases with over 430 mutations in 13 genetically distinct genes (*CLN1-8* and *CLN10-14*) [1]. Germline mutations in *CLN5*, resulting in either amino acid conversion (R112H, N192S, D279N) or truncations due to premature stop codons (W75X and Y392X), are causes of CLN5 disease. The truncation mutation, CLN5^Y392X^, is the most common mutation in patients. This form of NCL has an early onset between the ages of 3 - 7, with a lifespan into the teenage years. Patients exhibit symptoms including retinopathy, motor disorders, epilepsy, and cognitive regression. The principal characteristics leading to the identification of NCL disorder is aberrant lysosomal function and excess accumulation of auto fluorescent ceroid lipopigments in neurons and peripheral tissues [2].

CLN5 is a glycosylated endolysosomal protein that is translated as a type II integral membrane protein (residues 1 - 407), prior to being cleaved into a mature soluble form (93 - 407), which is then transported to endolysosomes [3]. Recent work has suggested that CLN5 could function as a glycoside hydrolase, but endogenous targets have not been identified [4]. Furthermore, CLN5 has been linked to the regulation of lysosomal pH [5] and mitophagy [6]. To further our understanding of this protein, using small interfering RNA (siRNA), we generated CLN5 knockdown (CLN5^KD^) HeLa cells. In these cells, we found less membrane bound Rab7A [7]. Rab7A is a small GTPases that can bind to and recruit retromer, a protein complex that mediates endosome-to-trans Golgi Network (TGN) trafficking. Once recruited, retromer interacts with the cytosolic tails of the lysosomal sorting receptors cationic independent mannose 6- phosphate receptor (CI-MPR) and sortilin to drive their endosome-to-TGN retrieval [8-10]. In CLN5^KD^ HeLa cells, the lack of Rab7A recruitment resulted in the lysosomal degradation of CI-MPR and sortilin, and misrouting of the lysosomal enzyme cathepsin D, due to lack of retromer recruitment [7]. Beyond retromer recruitment, Rab7A also regulates the degradation of endocytic cargo such as the epidermal growth factor (EGF) receptor (EGFR) [11] by interacting with PLEKHM1 [12], regulates organelle movement by interacting with RILP [13, 14], and regulates the late stages of autophagy by mediating autophagosome/lysosome fusion through HOPS and other tethering factors [15].

Recently, we have shown that CLN3 is also required in the endosome-to-TGN retrieval of the lysosomal sorting receptors [16]. Mutations in the gene encoding for CLN3 results in the juvenile variant of NCL (JNCL), commonly known as Batten disease. JNCL is the most common cause of childhood dementia, with an age of onset between 5 to 10 years [17]. CLN3 is a protein of 438 amino acids with six transmembrane domains, whose N- and C- terminal ends are located in the cytosol. This glycosylated protein localizes to endolysosomal membranes among other intracellular locations [18, 19], and can interact with other CLN proteins including CLN5. However, the molecular function of these interactions are not known [20]. Unlike depletion of CLN5, deletion of CLN3 did not affect the localization or activation of Rab7A or its effector retromer. Rather, CLN3 is required for the efficient Rab7A interaction with retromer, and for the retromer/sortilin interaction, most likely acting as a scaffold protein [16].

In this study, we generated CLN5 knockout (CLN5^KO^) HeLa cells on the same parental cell line used to generate CLN3 knockout (CLN3^KO^) and Rab7A knockout (Rab7A^KO^) cells [16]. This knockout system has enabled us to study the effects of a CLN5 disease-causing mutation. We extended our studies to identify defects in endocytic degradation and autophagy. Importantly, we identified significant decreases in previously known interactions between CLN3 and sortilin, retromer and Rab7A in CLN5^KO^ cells compared to wild-type cells. This leads to decreases in the Rab7A/retromer and retromer/sortilin interactions. Overall, our data suggest that CLN3 and CLN5 form a late endolysosome complex that modulates a subset of Rab7A functions. These finding provides insights into the pathogenic mechanisms observed in CLN3 and CLN5 disease.

## Materials and Methods

### Plasmids and Mutagenesis

RlucII-CLN3, GFP10-CLN3, Vps26A-nLuc, PLEKHM1-GFP10, RlucII-Rab7A, Vps26A- GFP10, RILP-GFP10 and sortilin-myc were previously described [16, 21, 22]. The various CLN5 mutants were engineered using site-directed mutagenesis from the previously described HA- CLN5 and CLN5-HA constructs [3]. Sortilin-YFP was a generous gift from Dr. Makoto Kanzaki, Tohoku University. mCherry-LC3 was a generous gift from Dr. Peter K. Kim, Sickkids Hospital. Lamp1-GFP was a generous gift from Dr. Juan Bonifacino, NICHD, NIH. mTagRFP-mWasabi- LC3 was a generous gift from Jian Lin, Peking University.

### Antibodies

The following mouse monoclonal antibodies were used: anti-actin (Wb: 1:3000, BD Biosciences 612657); anti-Lamp2 (Wb: 1:500, Abcam ab25631); anti-HA (Wb: 1:1000, Cedarlane Labs 901503); anti-myc (Wb: 1:1000, ThermoFisher Scientific LS132500), anti-Cathepsin D (Wb: 1:100, Sigma-Aldrich, IM03); anti-Flag (Wb: 1:1000, Sigma-Aldrich F1804). The following rabbit monoclonal antibodies were used: anti-Rab7A (Wb: 1:1000, Cell Signalling Technology D95F2); anti-EGFR (Wb: 1:1000, Abcam ab52894); anti-CLN5 (Wb: 1:1000 Abcam ab1700899). The following rabbit polyclonal antibodies were used: anti-Vps26A (Wb: 1:2000, Abcam ab23892); anti-FLAG (Wb: 1:1000, BioLegend, 902401); anti-sortilin (Wb: 1 µg/ml, Abcam ab16640).

### Cell culture and transient transfections

HeLa cells were cultured in Dulbecco’s modified Eagle’s medium (DMEM) supplemented with 2mM L-Glutamine, 100 U/ml penicillin, 100g/ml streptomycin and 10% FBS (Wisent Bioproducts, St-Bruno, QC) at 37 °C in a humidified chamber at 95% air and 5% CO_2_. Cells were seeded at a density of 2 × 10^5^/well for 12 well plates and 5 × 10^5^/well for 6 well plates 24 hours prior to transfection. Transfections were performed with polyethylenimine (PEI) (ThermoFisher Scientific, Ottawa, ON). Briefly, solution 1 was prepared by diluting plasmid into Opti-MEM (ThermoFisher Scientific). Solution 2 was prepared by diluting PEI (1 µg/µl) into Opti-MEM in a ration of 1:3 with the DNA to be transfected. After a 5 min incubation, the two solutions were mixed, vortexed for 3 s, incubated at room temperature (RT) for 15 min and subsequently added to the cells.

### CRISPR/Cas9 Editing

HeLa cells were transfected with an all-in-one CRISPR/Cas9 plasmid for CLN5 (plasmid number HCP202087-CG01-1-B, Genecopoeia, Rockville, MD). 72 hours post-transfection, cells were treated with 1mg/ml Geneticin (ThermoFisher Scientific) for 1 week. Limited dilution was performed to isolate single cells, which were allowed to grow for 2 weeks. Western blotting was used to identify CLN5^KO^ cells. Genomic DNA extracted form CLN5^KO^ HeLa cells was sequenced to confirm CLN5^KO^ cells. Rab7A^KO^ HeLa cells were previously described [16].

### BRET titration experiments

HeLa cells were seeded in 12-well plates and transfected with the indicated plasmids. 48 hours post transfection, cells were washed in PBS, detached with 5mM EDTA in PBS and collected in 500 µl of PBS. Cells were transferred to opaque 96-well plates (VWR Canada, Mississauga, ON) in triplicates. Total fluorescence was first measured with the Tecan Infinite M1000 Pro plate reader (Tecan Group Ltd., Mannedorf, Switzerland) with the excitation and emission set at 400 nm and 510 nm respectively for BRET^2^ and 500 nm and 530 nm for BRET^1^. The BRET^2^ substrate coelenterazine 400a and BRET^1^ substrate h-coelenterazine were then added to all wells (5 µM final concentration) and the BRET^2^ or BRET^1^ signals measured 2 min later. The BRET signals were calculated as a ratio of the light emitted at 525 ± 15 nm over the light emitted at 410 ± 40 nm. The BRET_net_ signals were calculated as the difference between the BRET signal in cells expressing both fluorescence and luminescence constructs and the BRET signal from cells where only the luminescence fused construct was expressed.

### Western blotting

Cell were detached using 5mM EDTA in PBS, washed in 1X PBS and collected by centrifugation. TNE buffer (150 mM NaCl, 50 mM Tris, pH 7.5, 2 mM EDTA, 0.5% Triton X-100 and protease inhibitor cocktail) was used to lyse cells by incubating them for 30 minutes on ice. Lysates were centrifuged at high speed for 10 minutes and the supernatants (whole cell lysate) were collected to be analyzed by Western blotting. Samples were mixed with sample buffer 3X to obtain a final concentration of 1X (62.5 mM Tris-HCl pH 6.5, 2.5% SDS, 10% glycerol, 0.01% bromophenol blue). Samples were incubated at 95°C for 5 minutes and resolved on SDS-PAGE followed by wet-transfer to nitrocellulose membranes. Detection was done by immunoblotting using the indicated antibodies.

### Membrane separation assay

24hrs after transfection, cells were collected in 5mM EDTA in PBS. The cells were subsequently snap frozen in liquid nitrogen and allowed to thaw at RT for 5min. The cells were resuspended in Buffer 1 (0.1 M Mes-NaOH pH 6.5, 1 mM MgAc, 0.5 mM EGTA, 200 M sodium orthovanadate, 0.2 M sucrose) and centrifuged at 10,000 g for 5 minutes at 4°C. The supernatants containing the cytosolic proteins (S, soluble fraction) were collected. The remaining pellet was resuspended in Buffer 2 (50 mM Tris, 150 mM NaCl, 1 mM EDTA, 0.1% SDS, 1% Triton X-100) and centrifuged at 10,000 g for 5 minutes at 4°C. Samples were loaded into SDS-PAGE gels in equal volumes. Fiji was used to quantify the intensity of the bands [23]. The intensity of each fraction was calculated and divided by the total intensity to determine the distribution of proteins.

### Cycloheximide chase

Wild-type, CLN5^KO^, Rab7A^KO^ HeLa cells were seeded in 6-well plates the day prior to transfection. 500 ng of HA fused wild-type and mutation harbouring CLN5 were transfected into CLN5^KO^ HeLa cells. 48 hours after transfection, cells were treated with 50 µg/ml of cycloheximide in Opti-MEM. Lysates were collected at 0, 3 or 6 hour-time points and run on 10% SDS-PAGE gels. Fiji was used to quantify the intensity of the bands [23]. All protein levels were standardized to the actin loading control. Amount of remaining protein is expressed as a percentage of the 0 time point in each group.

### EGFR degradation assay

Wild-type, CLN5^KO^ and Rab7A^KO^ cells were seeded in 6-well plates the day before the assay. In order to prevent *de novo* synthesis EGFR during EGF stimulation, cells were treated with 50 µg/ml cycloheximide in Opti-MEM for 1 hour. Stimulation of EGF was performed with 100 ng/ml EGF in Opti-MEM containing 50 µg/ml of cycloheximide. Cell lysates were collected as indicated above and Western blotting was performed. Fiji was used to quantify the intensity of the bands [23]. All protein levels were standardized to the actin loading control. Amount of remaining protein is expressed as a percentage of the 0 time point in each group.

### EGF-488 pulse-chase

Wild-type and CLN5^KO^ HeLa cells were seeded on coverslips the day before the experiment. Cells were serum-starved in Opti-Mem for 1 h followed by a 30-min pulse of 300 ng/ ml of EGF-488. Cells were then washed with PBS, fixed in 4% paraformaldehyde at 0 and 30 min and mounted onto slide using Fluoromount-G (ThermoFisher Scientific). The coverslips were sealed using nail polish. Cells were imaged using fluorescence microscope using a 63X objective. The number of puncta per cell was counted using Fiji. Briefly, the images were split into their individual channels. Using the green channel, the threshold was adjusted to 0,93% in order to get signal only from the puncta. The number of puncta counted from an image was divided by the number of nuclei within that image, which gave us the number of puncta per cell. 35 cells per condition for each time point were analyzed.

### Acyl-RAC to isolate palmitoylated proteins

The isolation of palmitoylated protein was adapted from Modica et al., 2017. Briefly, cells were lysed in TNE (150 mM NaCl, 50 mM Tris, pH 7.5, 2 mM EDTA, 0.5% Triton X-100 and protease inhibitor cocktail) supplemented with 50mM N-Ethylmaleimide (NEM) and incubated 30 minutes in rotating wheel at 4°C. Samples were centrifuged 10 minutes at 10000g at 4°C and the collected supernatants were incubated 2 hours at RT on a rotating wheel. Samples were then precipitated over night with two volumes of cold acetone at −20°C to remove NEM. After washing with cold acetone, the pellets were resuspended in binding buffer (100 mM HEPES, 1 mM EDTA, 1% SDS) with 250 mM hydroxylamine (Sigma-Aldrich) (NH2OH) pH7.5 to cleave palmitate residues off of proteins. Control samples were resuspended in binding buffer containing 250mM NaCl. When the pellets were completely resuspended, water-swollen thiopropyl sepharose 6B beads (GE Healthcare Life Sciences, Mississauga, ON) were added and samples were incubated 2 hrs at RT on rotating wheel. Beads were then washed 4 times with binding buffer and captured proteins were eluted with sample buffer containing 100mM DTT.

### Autophagy assays

Cells were seeded on 6-well plates containing glass coverslips. The next day, the cells were transfected with the indicated plasmids. 48 hours post-transfection cells, the cells were starved in EBSS for 3 hours. Then, the coverslips were washed with PBS and fixed with 4% paraformaldehyde in PBS for 12 min at room temperature and washed twice with PBS. Fixed cells on the coverslips are mounted on cover slides with Fluoromount G (Thermo Fisher Scientific). Images were acquired using a fluorescence microscope using a 63X objective. Quantification of the fluorescence from 35 cells to analyze the colocalization of LC3 and Lamp1 was performed using Fiji software by splitting the red and green channels and manually outlining the cells to obtain the region of interest (ROI). Coloc2 was performed to determine the Pearson’s coefficient. Red (LC3) in green (Lamp1) was chosen together with the bisection option for the threshold regression. For autophagic flux comparison, intensity of the red signal is divided by the green signal for each cell. Quantification of the fluorescence from 30 cells to analyze the quenching of the green signal from the mTagRFP-mWasabi-LC3 probe was performed using the Fiji software. Red and green channels were split, and the cells manually outlined to get the ROI. Red and green fluorescence total intensities coming from each selected cells are analyzed.

## Results

### Retromer is not efficiently recruited to endosomal membranes in CLN5^KO^ HeLa cells

In our previous work conducted in CLN5^KD^ HeLa cells, we showed decreased membrane recruitment of Rab7A leading to a significant decrease in the membrane recruitment of its effector, retromer [7]. In order to study the role of CLN5 disease-causing mutations, we generated CLN5^KO^ HeLa cells using CRISPR/Cas9. Our sequencing data and Western blot analysis confirm the deletion of CLN5 in this cell line (**Fig. S1A and B)**. To confirm our previous results obtained in CLN5^KD^ cells, we performed a membrane isolation experiment in our CLN5^KO^ HeLa cells as we have previously done [7, 16, 22], and compared the membrane distribution of Rab7A in wild-type and CLN5^KO^ HeLa cells (**Fig. 1A**). Our membrane separation was successful, as the integral membrane protein Lamp2 was found in the pellet fraction (P) containing membranes, and not in the supernatant fraction (S) contains the cytosol (**Fig. 1A**). Quantification of 3 independent experiments showed that CLN5 deletion had no significant effect on Rab7A membrane distribution (**Fig. 1B**). We sought to determine the effect of a CLN5 mutation known to cause human disease on Rab7A membrane recruitment. Thus, we engineered a CLN5 disease-causing mutation expression plasmid using site directed mutagenesis to generate CLN5^Y392X^-HA, and expressed it in CLN5^KO^ HeLa cells (**Fig. 1A**). Rab7A distribution was not affected in CLN5^KO^ HeLa cells expressing CLN5-HA or expressing the disease causing mutation, CLN5^Y392X^-HA (**Fig. 1B**). The small GTPase Rab7A regulates the spatiotemporal recruitment of retromer [9, 10]. Therefore, we repeated the membrane assay as above to test if CLN5 is required for membrane recruitment of retromer. This time we included Rab7A^KO^ HeLa cells as a control, as we previously demonstrated a reduction of retromer recruitment in these cells [16]. Compared to wild-type HeLa cells, we observed a significant decrease in retromer recruitment (as detected by the retromer subunit Vps26A) in CLN5^KO^ and Rab7A^KO^ HeLa cells (**Fig. 1A**). Quantification of 6 independent experiments found a reduction of 21.4% in the membrane recruitment of retromer in CLN5^KO^, compared to a 31.4% decrease in recruitment in Rab7A^KO^ HeLa cells (**Fig. 1C**). These values are comparable to previous studies [9, 10]. This decrease in retromer membrane recruitment was rescued by expressing CLN5-HA in CLN5^KO^ cells, indicating the effect seen on Vps26A recruitment is specific to CLN5 deletion, and likely not an off-target effect (**Fig. 1A and C**). The expression of CLN5^Y392X^-HA in CLN5^KO^ cells partially rescued retromer recruitment, but not to levels comparable as that of wild-type CLN5 (**Fig. 1C**).

**Figure 1.**
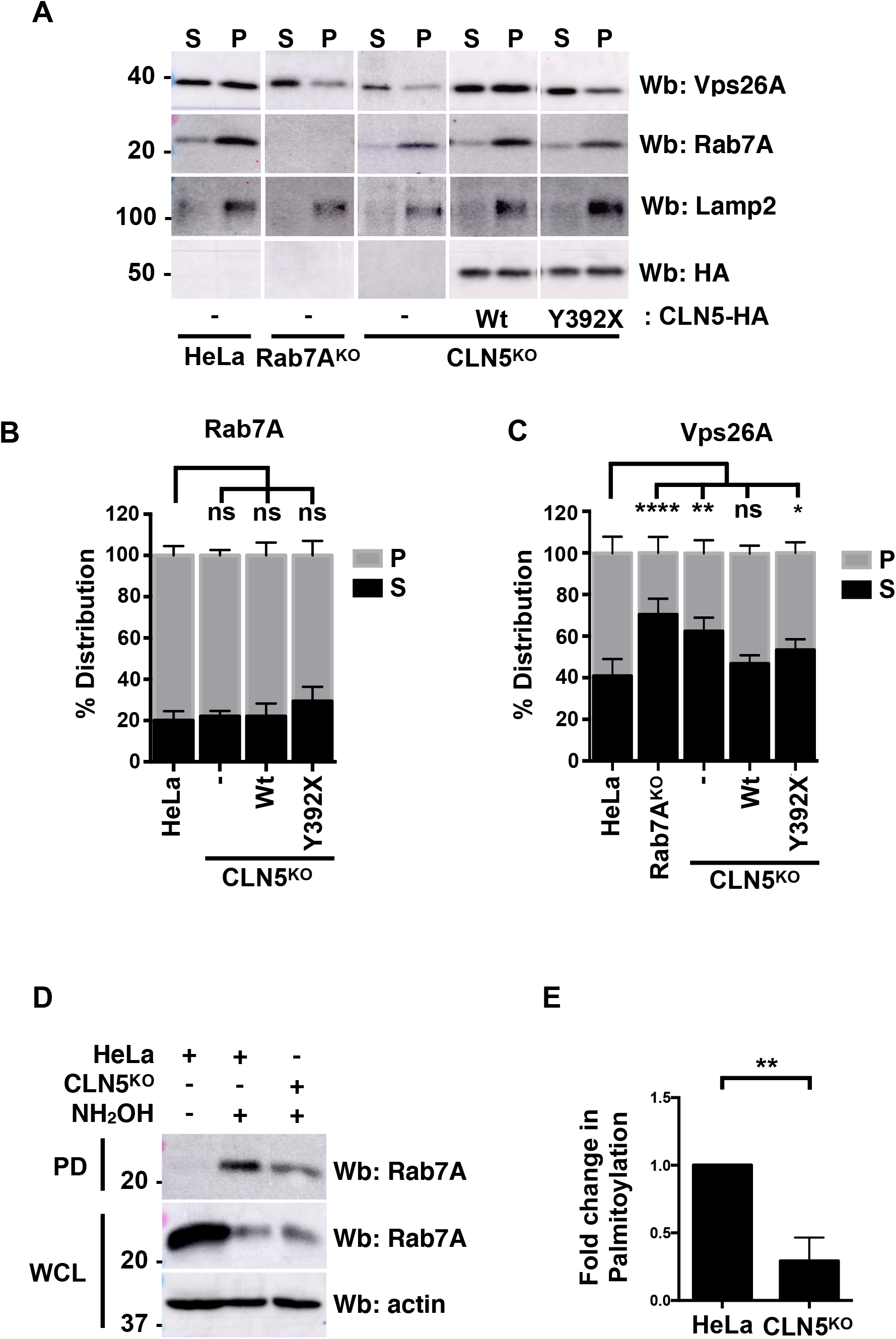
Rab7A palmitoylation is reduced in CLN5^KO^ HeLa cells affecting retromer recruitment. (**A**) A membrane separation assay was performed on wild-type, Rab7A^KO^, CLN5^KO^, and CLN5^KO^ HeLa cells expressing CLN5-HA or CLN5^Y392X^-HA. Western blotting (Wb) was performed with anti-Rab7A, anti-Vps26A, anti-Lamp2 (as a marker of the membrane fraction) and anti-HA (to identify cells expressing CLN5-HA) antibodies. S; supernatant fraction, P; pellet fraction. (**B**) Quantification of 3 separate membrane isolation assay experiments for Rab7A distribution. Data is shown as mean±s.d.; ns, not significant; two-way ANOVA followed by Tukey’s post hoc test. (**C**) Quantification of 3 separate membrane isolation assay experiments for Vps26A distribution. Data is shown as mean±s.d.; ns, not significant; *P≤0.01, **P≤0.01,****P≤0.0001, two-way ANOVA followed by Tukey’s post hoc test (**D**) Whole cell lysate (WCL) from wild-type and CLN5^KO^ HeLa cells were subjected to Acyl-RAC analysis to determine the palmitoylation status of Rab7A. NH_2_OH: hydroxylamine, PD: pull-down (**E**) Quantification of 3 separate Acyl-RAC assay experiments. Data is shown as mean±s.d.; **P≤0.01, Student’s *t*-test.

The question remained as to how Rab7A was membrane bound, but not able to recruit retromer. In recent years, Rab7A has been shown to be phosphorylated [24-26], ubiquitinated [27] and palmitoylated [22]. These post-translational modifications have been demonstrated to regulate various functions of this small GTPase, often by modulating the interaction with its effectors. In particular, we have previously shown that Rab7A palmitoylation is required for the recruitment of retromer to endosomal membranes. While non-palmitoylatable Rab7A is still membrane bound, it does not efficiently interact with retromer, and is not capable of rescuing retromer membrane recruitment in Rab7A^KO^ HEK293 cells [22]. Since Rab7A was still membrane bound in CLN5^KO^ HeLa cells, but retromer was not, we used Acyl-RAC to test if Rab7A palmitoylation is affected in these cells (**Fig. 1D**). We found that Rab7A palmitoylation is significantly decreased in CLN5^KO^ HeLa cells compared to wild-type cells (**Fig. 1E**).

### CLN5 is required for efficient retromer interactions

Rab7A coordinates the spatiotemporal recruitment of retromer to endosomal membranes [9, 10]. Since we observed decreased retromer membrane recruitment along with decreased Rab7A palmitoylation, we expected to observe a weaker interaction between Rab7A and retromer in CLN5^KO^ HeLa cells. To test our hypothesis, we used bioluminescence resonance energy transfer (BRET), as we have previously done [16, 22]. Compared to co-immunoprecipitation, BRET is performed in live cells, with proteins localized to their native environment. From BRET titration curves, the BRET^50^ can be calculated, which is the value at which the concentration of the acceptor is required to obtain 50% of the maximal BRET signal (BRET_max_). The BRET_50_ is indicative of the propensity of a pair of protein to interact, as the smaller the BRET_50_, the stronger the interaction [28, 29]. The energy donor Renilla Luciferase II (RlucII) was fused at the N-terminus to wild-type Rab7A (RlucII-Rab7A), while the energy acceptor GFP10 was fused at the C-terminus of the retromer subunit Vps26A. Neither of these tags appear to interfere with the function of the proteins, as we have previously shown that RlucII had little effect on the distribution or function of Rab7A [22], while we confirmed that Vps26A-GFP10 is efficiently integrated into the retromer trimer [16]. We have previously used BRET to study the Rab7A/Vps26A interaction [22]. This interaction is specific as Rab7A did not interact with AP-1 subunits (a clathrin adaptor that has been localized to both the TGN and endosomes), while Vps26A did not interact with RlucII-Rab1a, which is localized to the Golgi apparatus [22]. Wild-type and CLN5^KO^ HeLa cells were co-transfected with a constant amount of RlucII-Rab7A, and increasing amounts of Vps26A-GFP10 to generate BRET titration curves (**Fig. 2A, blue and red curves**). The BRET signal between RlucII-Rab7A and Vps26A-GFP10 rapidly increased with increasing amounts of expressed Vps26A-GFP10 until it reached saturation, suggesting a specific interaction. Compared to wild-type HeLa cells, we found a 5- fold increase in the BRET_50_ value for Rab7A binding to Vps26A in CLN5^KO^ cells, suggesting a weaker interaction (**Fig. 2B**). To test if the disease-causing mutation in CLN5 affected the Rab7A/retromer interaction, we expressed wild-type HA-CLN5 (**Figure 2A, green curve**) or HA-CLN5^Y392X^ (**Figure 2A, purple curve**) in CLN5^KO^ cells and generated BRET titration curves as above. While expressing HA-CLN5 rescued the Rab7/retromer interaction as the BRET_50_ was similar to the BRET_50_ in wild-type HeLa cells (**Fig. 2B**), expression of HA-CLN5^Y392X^ did not, as the BRET_50_ value was 4 fold larger (**Fig. 2B**). Our previous work suggests that palmitoylation favours the association of Rab7A to a specific endosomal domain to optimize its interaction with retromer [22], without affecting the overall ability of the small GTPase to interact with this effector. As such, when the Rab7A/retromer interaction was analyzed with BRET, hence preserving cellular membranes, we observed a decreased interaction, while no change in the interaction was observed in co-immunoprecipitation where cellular compartments are lost. To confirm if this was the case in CLN5^KO^ HeLa cells, we performed co-immunoprecipitation between Rab7A and retromer in wild-type and CLN5^KO^ HeLa cells (**Fig. S1C**). We found no change in the Rab7A/retromer interaction using this method.

**Figure 2.**
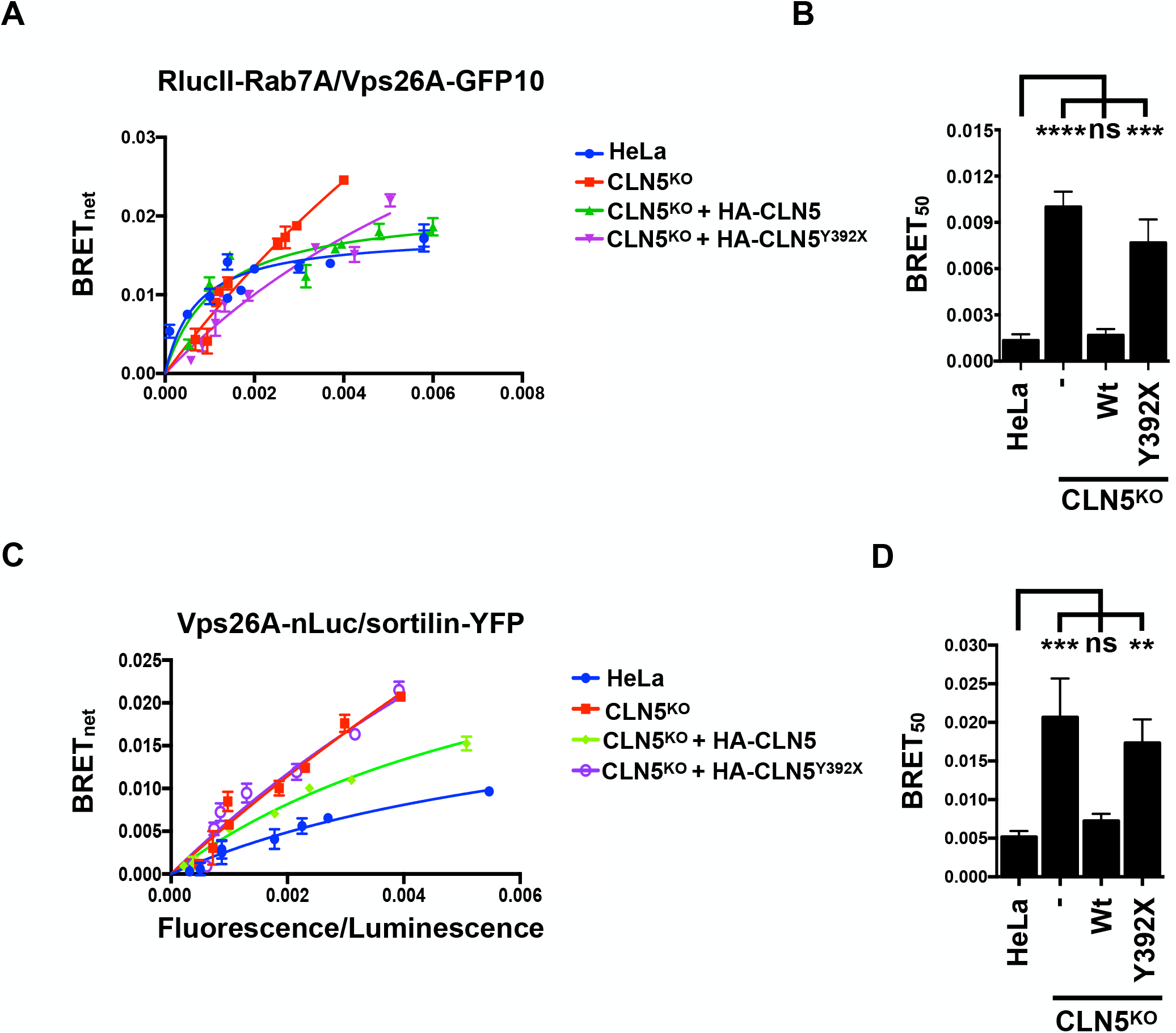
Weaker Rab7A/retromer and retromer/sortilin interactions in CLN5^KO^ HeLa cells. (**A**) Wild-type, CLN5^KO^ and CLN5^KO^ HeLa rescued with wild-type HA-CLN5 or HA-CLN5^Y392X^ were transfected with a constant amount of RlucII-Rab7A and increasing amounts of Vps26A- GFP10 to generate BRET titration curves. BRET signals are plotted as a function of the ratio between the GFP10 fluorescence over RlucII luminescence. (**B**) BRET_50_ was extrapolated from 3 independent experiments. Data is shown as mean±s.d.; ns, not significant, ***P≤0.001, ****P≤0.0001, one-way ANOVA followed by Tukey’s post hoc test (**C**) Wild-type, CLN5^KO^ and CLN5^KO^ HeLa rescued with wild-type HA-CLN5 or HA-CLN5^Y392X^ were transfected with a constant amount of Vps26A-nLuc and increasing amounts of sortilin-YFP to generate BRET titration curves. BRET signals are plotted as a function of the ratio between the YFP fluorescence over nLuc luminescence. (**D**) Quantification of 3 independent experiments. Data is shown as mean±s.d.; ns, not significant; **P≤0.01, ***P≤0.001, one-way ANOVA followed by Tukey’s post hoc test

CI-MPR and sortilin are known to interact with retromer, which is necessary for their endosome-to-TGN trafficking [8, 30, 31]. Using BRET, we have previously shown that nanoLuciferase-tagged Vps26A (Vps26A-nLuc) and YFP-tagged sortilin (sortilin-YFP) interact, and that this interaction is significantly weaker in CLN3^KO^ cells compared to wild-type HeLa cells [16]. We tested the sortilin/retromer interaction in CLN5^KO^ HeLa cells to determine if this protein plays a role in this interaction. Wild-type and CLN5^KO^ HeLa cells were co-transfected with a constant amount of Vps26A-nLuc and increasing amounts of sortilin-YFP to generate BRET titration curves (**Fig. 2C, blue and red curves**). We found a significantly weakened interaction between retromer and sortilin in CLN5^KO^ cells compared to wild-type HeLa cells, as we observed a 4 fold increase in the BRET_50_ value (**Fig. 2D**). The retromer/sortilin interaction in CLN5^KO^ HeLa cells could be rescued by expressing HA-CLN5 (**Fig. 2C, green curve**), as the BRET_50_ value was similar to the one we calculated in wild-type HeLa cells. We next performed BRET experiments to determine the impact of the CLN5^Y392X^ disease-causing mutation on the Vps26A/sortilin interaction. Expression of HA-CLN5^Y392X^ in CLN5^KO^ HeLa cells (**Fig. 2C, purple curve**) did not rescue the interaction as shown by the BRET_50_ value (**Fig. 2D**). To confirm our BRET data, we immunoprecipitated endogenous Vps26A with anti-Vps26A antibody, and blotted for endogenous sortilin (**Fig. S1C**). By co-immunoprecipitation, we observed no change in the Vps26A/sortilin interaction in CLN5^KO^ cells, suggesting that the proteins can interact when the contribution of membrane distribution is not considered.

### Sortilin is degraded in CLN5^KO^ HeLa cells

In cells lacking functional retromer, sortilin does not efficiently recycle to the TGN, accumulates in late endosomes, and is subsequently degraded in lysosomes [16]. We have previously demonstrated the same phenotype in cells depleted of CLN5 [7]. Using our CLN5^KO^ HeLa cells, we performed a cycloheximide chase experiment to determine receptor stability as we have previously done [7, 16, 32]. Wild-type, Rab7^KO^ and CLN5^KO^ HeLa cells were incubated with serum free medium containing 50 µg/ml cycloheximide and collected after 0, 3, and 6 hours of incubation. Western blotting was performed using anti-sortilin antibody, anti-actin antibody (as a loading control), while HA staining was used to demonstrate the expression level of HA tagged CLN5 constructs. Western blot (Wb) analysis shows decreased levels of sortilin in CLN5^KO^ HeLa cells at 3 and 6 hours compared to Rab7^KO^ and wild-type HeLa cells (**Fig. 3A**). Transfecting CLN5-HA in CLN5^KO^ cells rescued this phenotype (**Fig. 3A**). We next aimed to determine the impact of CLN5^Y392X^ on the stability of sortilin. Expression of CLN5^Y392X^-HA in CLN5^KO^ cells did not rescue the degradation of sortilin (**Fig. 3A**). Quantitative analysis of 3 independent cycloheximide chase experiments showed that compared to wild-type cells which had 76.6% and 74.75% of sortilin remaining at 3 and 6 hours respectively (**Fig. 3B**), sortilin was significantly degraded in CLN5^KO^ cells, as only 23.8% and 22.8% of sortilin remained after 3 and 6 hours (**Fig. 3B**). As expected, we observed no significant degradation in Rab7A^KO^ cells, which had 75.7% and 84.3% sortilin remaining at 3 and 6 hours respectively (**Fig. 3B**). Expressing CLN5-HA in CLN5^KO^ cells rescued the stability and hence recycling of sortilin, as it was no longer degraded and had protein levels remaining of 79.6% and 77% respectively at 3 and 6 hours, which is similar to wild-type HeLa cells (**Fig. 3B**). We observed a partial rescue of sortilin degradation in CLN5^KO^ cells expressing CLN5^Y392X^-HA at the 3 hours time point, as 38.4% was remaining. However, after 6 hours, only 17.7% was remaining, suggesting that this disease-causing mutation did not rescue the knockout phenotype.

**Figure 3.**
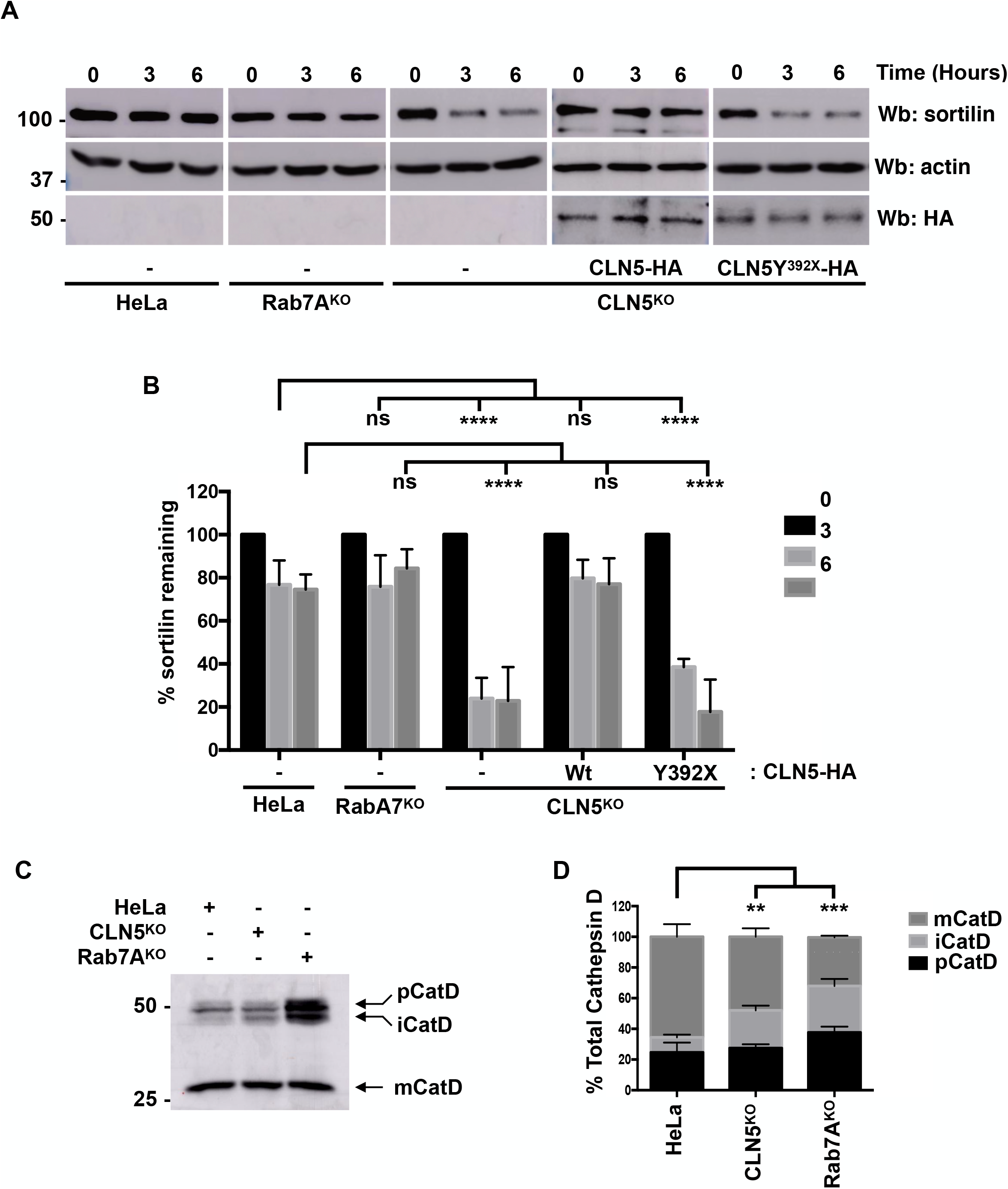
The lysosomal sorting receptor sortilin is degraded in CLN5^KO^ HeLa cells. (**A**) Wild-type, Rab7A^KO^, CLN5^KO^, and CLN5^KO^ HeLa cells expressing CLN5-HA or CLN5^Y392X^- HA were treated with 50 µg/ml of cycloheximide in serum free media for the indicated times. Whole cell lysate was run on a SDS-PAGE and Western blotting (Wb) was performed with anti-sortilin, anti-actin and anti-HA antibodies. (**B**) Quantification of 3 independent experiments. Data is shown as mean±s.d.; ns, not significant; ****P≤0.0001, two-way ANOVA followed by Tukey’s post hoc test. (**C**) Whole cell lysate from wild-type, CLN5KO and Rab7AKO HeLa cells was resolved by SDS-PAGE and Western blotting (Wb) was performed using anti-cathepsin D antibody. (**D**) Quantification of pro-cathepsin D (proCatD, 53 kDa), intermediate cathepsin D (iCatD, 48 kDa) and mature cathepsin D (mCatD, 31 kDa). The amount of each is expressed as a percentage of total cathepsin D. Statistical significance refers to mature Cathepsin D (mCatD). Data are mean±s.d.; ns, not significant; **P≤0.01,***P≤0.001, two-way ANOVA followed by Tukey’s post hoc test.

Since we observed the degradation of sortilin in CLN5^KO^ HeLa cells, we aimed to determine if this had an impact on lysosomal function. Cathepsin D is generated as a 53 kDa pro-cathepsin D (proCatD), which is processed to a 47 kDa intermediate form (iCatD), before being fully processed to a 31 kDa mature cathepsin D (mCatD) once it reaches the lysosome [33]. Depletion of Rab7A by siRNA affects this processing, resulting in increased amounts of the pro and intermediate forms [9]. Total cell lysate from wild-type, CLN5^KO^, and Rab7A^KO^ HeLa cells was run on a SDS-PAGE and Western blotting was performed using anti-cathepsin D antibody (**Fig. 3C**). Compared to wild-type HeLa cells which had 24.6% pro-cathepsin (proCatD), 9.8% intermediate (iCatD) and 65.6% mature cathepsin D (mCatD), CLN5^KO^ HeLa had significantly increased iCatD (24.6%) and significantly reduced mCatD (48%), with similar levels of proCatD (27.4%). On the other hand, Rab7A^KO^ HeLa cells had increased levels of proCatD (37.6%) and iCatD (30.3%), and had decreased amounts of mCatD (32.1%) (**Fig. 3D**).

### CLN5 is required for efficient CLN3 interactions

Recently, we showed that CLN3 functions as a scaffold protein to ensure the Rab7A/ retromer and retromer/sortilin interactions, which are sequentially required to regulate the endosome-to-TGN retrieval of sortilin [16]. In this current study, we demonstrated a role for CLN5 in regulating the stability of sortilin. However, the question remains how a soluble lysosomal protein, CLN5, can regulate the Rab7A/retromer and retromer/sortilin interactions, which occur in the cytosol. In order to better understand the relationship of CLN5 and CLN3 in this trafficking pathway, we wanted to determine the role of CLN5 in modulating CLN3 interactions. Previous studies have shown that CLN3 and CLN5 interact [20, 34]. We confirmed the CLN3/CLN5 interaction using co-immunoprecipitation, and tested the impact of a disease-causing mutation in CLN5 on this interaction (**Fig. S2A**). The disease-causing mutation we tested, CLN5^Y392X^, had no impact on the CLN3/CLN5 interaction (**Fig. S2A**). The CLN3/Rab7A interaction has been previously shown using co-immunoprecipitation and BRET experiments [16, 35]. To determine if CLN5 plays a role in this interaction, wild-type (**Fig. 4A, blue curve**) and CLN5^KO^ HeLa cells (**Fig. 4A, red curve**) were co-transfected with a constant amount of RlucII-Rab7A and increasing amounts of GFP10-CLN3 to generate BRET titration curves. We extrapolated the BRET_50_ for the interaction between CLN3 and Rab7A and found that the BRET_50_ was 2.85 fold larger in CLN5^KO^ HeLa cells compared to wild-type cells, indicating a weaker CLN3/Rab7A interaction in cells lacking CLN5 (**Fig. 4B**). Expressing HA-CLN5 in CLN5^KO^ cells rescued the CLN3/Rab7A interaction (**Fig. 4A, green curve**) as the BRET_50_ was similar to wild-type HeLa cells (**Fig. 4B**). Expressing HA-CLN5^Y392X^ in CLN5^KO^ cells had a partial rescue effect on the CLN3/Rab7A interaction (**Fig. 4A, purple curve**), but the BRET_50_ was still more than 2 fold larger (**Fig. 4B**).

**Figure 4.**
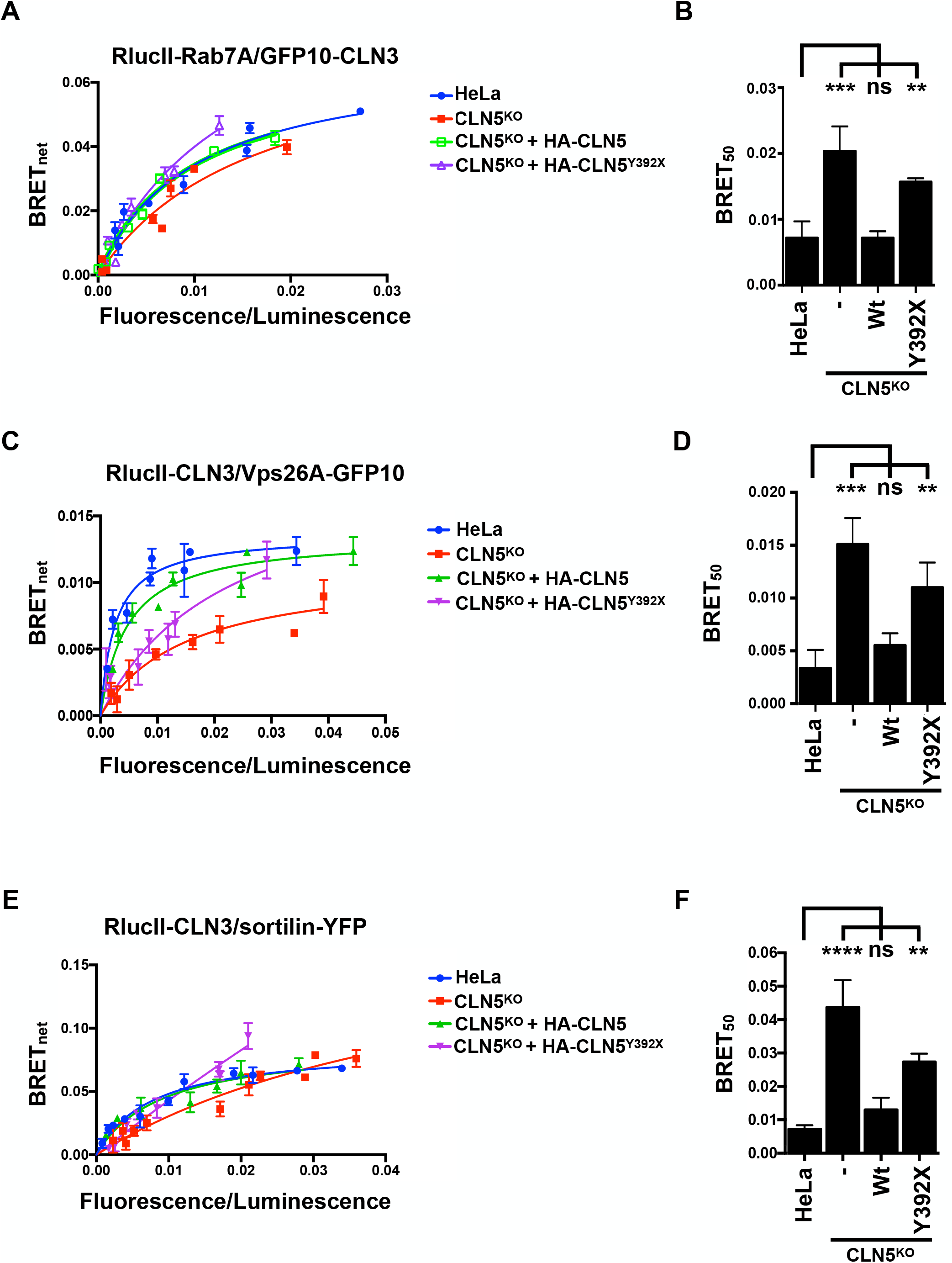
CLN5 modulates CLN3 interactions. (**A**) Wild-type, CLN5^KO^ and CLN5^KO^ HeLa rescued with wild-type HA-CLN5 or HA-CLN5^Y392X^ were transfected with a constant amount of RlucII-Rab7A and increasing amounts of GFP10- CLN3 to generate BRET titration curves. BRET signals are plotted as a function of the ratio between the GFP10 fluorescence over RlucII luminescence. (**B**) BRET_50_ was extrapolated from 3 independent experiments. Data is shown as mean±s.d.; ns, not significant, **P≤0.01, ****P≤0.0001, one-way ANOVA followed by Tukey’s post hoc test. (**C**) Wild-type, CLN5^KO^ and CLN5^KO^ HeLa rescued with wild-type HA-CLN5 or HA-CLN5^Y392X^ were transfected with a constant amount of RlucII-CLN3 and increasing amounts of Vps26A-GFP10 to generate BRET titration curves. BRET signals are plotted as a function of the ratio between the GFP10 fluorescence over RlucII luminescence. (**D**) BRET_50_ was extrapolated from 3 independent experiments. Data is shown as mean±s.d.; ns, not significant, **P≤0.01, ***P≤0.0001, one-way ANOVA followed by Tukey’s post hoc test. (**E**) Wild-type, CLN5^KO^ and CLN5^KO^ HeLa rescued with wild-type HA-CLN5 or HA-CLN5^Y392X^ were transfected with a constant amount of RlucII- CLN3 and increasing amounts of sortilin-YFP to generate BRET titration curves. BRET signals are plotted as a function of the ratio between the YFP fluorescence over RlucII luminescence. (**F**) Quantification of 3 independent experiments. Data is shown as mean±s.d.; ns, not significant, **P≤0.01, ****P≤0.0001, one-way ANOVA followed by Tukey’s post hoc test

We previously demonstrated an interaction between CLN3 and retromer [16]. To test if CLN5 plays a role in modulating this interaction, we generated BRET titration curves using RlucII-CLN3 and Vps26A-GFP10 in wild-type (**Fig. 4C, blue curve**) and CLN5^KO^ HeLa cells (**Fig. 4C, red curve**). We found a significantly weaker CLN3/retromer interaction in CLN5^KO^ cells compared to wild-type HeLa cells as shown by the 4 fold increase of the BRET_50_ in CLN5^KO^ HeLa cells compared to wild-type cells (**Fig. 4D**). As expected, expressing HA-CLN5 in CLN5^KO^ cells rescued the CLN3/retromer interaction (**Fig. 4B, green curve**) as the BRET_50_ was similar to wild-type HeLA cells (**Fig. 4C**). Expressing HA-CLN5^Y392X^ in CLN5^KO^ HeLa cells had a partial rescue effect on the CLN3/retromer interaction (**Fig. 4C, purple curve**), but the BRET_50_ was still more than 3 fold larger (**Fig. 4D**).

We have also shown an interaction between CLN3 and sortilin [16] and between CLN5 and sortilin [7]. To test if a disease-causing mutation in CLN5 affected its ability to interact with sortilin, we performed a co-immunoprecipitation assay (**Fig. S2C**). Wild-type CLN5, as well as the disease-causing mutant tested, CLN5^Y392X^, were able to interact with sortilin. We next generated BRET titration curves using RlucII-CLN3 and sortilin-YFP in wild-type (**Fig. 4E, blue curve**) and CLN5^KO^HeLa cells (**Fig. 4E, red curve**). We extrapolated the BRET_50_for the interaction between CLN3 and sortilin and found that the BRET_50_ in CLN5^KO^ HeLa cells was 6 fold larger than in wild-type cells, indicating a decreased interaction in CLN5^KO^ cells (**Fig. 4F**). Expressing HA-CLN5 in CLN5^KO^ HeLa cells rescued the phenotype as the BRET_50_ value was similar to wild-type cells (**Fig. 4F, green curve**), while expressing HA-CLN5^Y392X^ (**Fig. 4F, purple curve**) did not. Although expression of CLN5^Y392X^ had a partial rescue of the CLN3/sortilin interaction, the BRET_50_ values were not restored to levels comparable to rescuing with wild-type CLN5 (**Fig. 4F**). To confirm our BRET results, we performed a co-immunoprecipitation experiment to determine the role of CLN5 in the modulating the CLN3/Rab7A, CLN3/retromer and sortilin/CLN3 interactions (**Fig. S2B**). We found that retromer (Vps26A), Rab7A and sortilin could co-immunoprecipitate with Flag-CLN3 in wild-type HeLa cells, but not in CLN5^KO^ HeLa cells. The interaction was specific as expressing empty Flag vector did not co-immunoprecipitate retromer or Rab7A (**Fig. S2B**).

### CLN5 is required for the efficient degradation of proteins following internalization

Another function of Rab7A is mediating key steps in the degradation of endocytic cargo [11, 36]. Upon Epidermal Growth Factor (EGF) stimulation, EGF receptor (EGFR) is internalized and can be either recycled to the cell surface, or degraded in lysosomes [37]. At least two Rab7A effectors have been implicated in EGFR degradation, RILP and PLEKHM1. Depletion of either of these proteins results in significant delays in the degradation kinetics of EGFR [12, 38, 39]. To determine whether CLN5 is implicated in this pathway, we investigated the degradation kinetics of EGFR in wild-type and CLN5^KO^ HeLa cells. The cells were serum starved for 1 hour in the presence of cycloheximide and then stimulated with 100 ng/ml of EGF in the presence of cycloheximide for the indicated periods of time. The level of endogenous EGFR was determined by Western blot using anti-EGFR antibody. Anti-actin staining was used as a loading control (**Fig. 5A**). In wild-type cells, EGFR degradation was observed after 15 minutes and quantification of 3 independent experiments found substantial degradation at 15 (34.3% remaining) and 120 minutes (9% remaining) (**Fig. 5B**). When we compared the degradation kinetics of EGFR in CLN5^KO^ cells, we found significant delay at 15 (75.6% remaining) compared to wild-type cells, with no significant difference at 120 minutes (10.3% remaining) (**Fig. 5B**).

**Figure 5.**
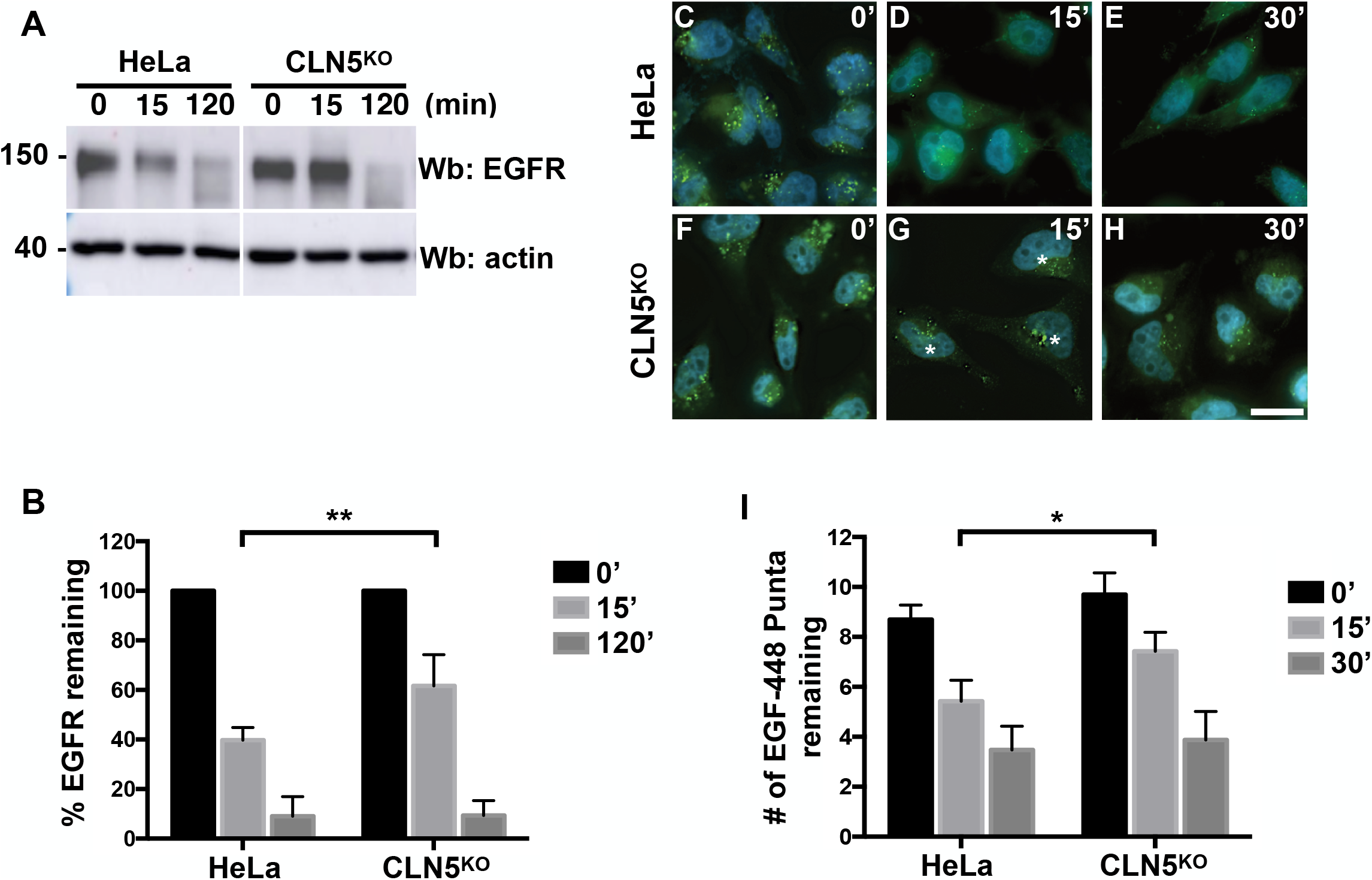
EGF and EGFR degradation is delayed in CLN5^KO^ cells. (**A**) Wild-type and CLN5^KO^ HeLa cells were incubated with 50 µg/ml cycloheximide for 1 h and subsequently treated with 100 ng/ml EGF in Opti-MEM for 0, 15 and 120 min. Whole cell lysate was then resolved by SDS-PAGE and a Western blot (Wb) was performed using anti-EGFR antibody. Anti-actin staining was used as a loading control. (**B**) Quantification of the remaining EGFR as detected in A was performed in 3 independent experiments. Data are mean±s.d. ns, not significant; **P≤0.01; one-way ANOVA followed by Tukey’s post hoc test. Scale bar: 10 µm. (**C - H**) Wild-type (**C - E**) and CLN5^KO^ (**F - H**) HeLa cells were grown on coverslips and incubated with 300 ng/ml EGF-488 for 0, 15 or 30 min. The cells were fixed with 4% PFA for 12 min, followed by staining with DAPI to visualize the nucleus. Images were taken on a Zeiss Fluorescence microscope using a 63× objective. Stars indicate remaining EGF-488. (**I**) EGF-488 (green puncta) were counted using ImageJ in 50 cells per condition. The results shown are the average number of puncta per cell per condition. Data are mean±s.d. ns, not significant; *P≤0.05; one-way ANOVA followed by Tukey’s post hoc test. Scale bar: 10 µm.

Next, we tested the degradation kinetics of Alexa-488 labeled EGF (EGF-488) using the same cell lines to confirm our EGFR degradation results. Following 2 hours of serum starvation, wild type (**Fig. 5C - E**) and CLN5^KO^ (**Fig. 5F - H**) HeLa cells were incubated with 300 ng/ml EGF-488 for 30 minutes, washed and then chased for 0 (**Fig. 5C and F**), 15 (**Fig. 5D and G**) or 30 minutes (**Fig. 5E and H**). Images were acquired at random from the different conditions and the number of EGF-488 puncta from 35 cells per condition were counted using Image J as we have previously done [16]. At time 0 min, both wild-type and CLN5^KO^ HeLa cells had comparable number of EGF-488 puncta (an average of 8.7 versus 9.7 respectively). After 15 minutes of chase, wild-type HeLa cells had an average of 5.4 puncta per cell, while CLN5^KO^ HeLa had on average 7.4 puncta per cell, an increase of 37% (**Fig. 5I**). After 30 minutes of chase, wild-type HeLa cells had an average of 3.4 puncta per cell, while CLN5^KO^ HeLa had on average 3.8 puncta per cell (**Fig. 5I**).

In order to understand the mechanism behind the delayed EGF and EGFR degradation, we used BRET to determine if the Rab7A/PLEKHM1 or Rab7A/RILP interactions were affected (**Fig. 6A - D**). RILP and PLEKHM1 are implicated in the degradation of internalized cargo such as EGFR by either participating in the tethering of vesicles, as is the case for PLEKHM1 [12], or by mediating the movement of vesicles, as is the case for RILP [40]. We generated BRET titration curves by transfecting a constant amount of RlucII-Rab7A and increasing amounts of PLEKHM1-GFP10 (**Fig. 6A**) or increasing amounts of RILP-GFP10 (**Fig. 6C**). We found no significant change in the interaction between PLEKHM1 and Rab7A in CLN5^KO^ HeLa cells compared to wild-type HeLa cells as shown by the similar BRET_50_ values (**Fig. 6B**). On the other hand, the interaction between RILP and Rab7A was significantly disrupted in CLN5^KO^ cells, as the BRET_50_ value was 3 fold higher for the Rab7A/RILP interaction in CLN5^KO^ cells compared to wild-type cells, suggesting a weaker interaction (**Fig. 6D**). Rab7A can interact with RILP to subsequently engage dynein motors for minus-end transport of lysosomes [14, 41]. If this transport system is affected due to the decreased Rab7A/RILP interaction, we would expect dysfunctional minus-end movement of lysosomes following starvation and the induction of autophagy. We compared the positioning of lysosomes, by staining with the lysosomal membrane protein CD63, in wild-type and CLN5^KO^ cells that were starved in EBSS for 3 hours or maintained in DMEM (**Fig. 6E - H**). In both wild-type (**Fig. 6E and E’**) and CLN5^KO^ (**Fig. 6F and F’**) HeLa cells cultured in DMEM, CD63 positive lysosomes were distributed throughout the cells. In nutrient starved wild-type cells, lysosomes efficiently moved in a retrograde manner towards the perinuclear region of the cell (**Fig. 6G and G’, white arrows**). However, while CLN5^KO^ HeLa cells were able to move some lysosomes, they were not as efficient as wild-type HeLa cells (**Fig. 6H and H’**). Quantification of 30 cells per condition showed that as a percentage of total CD63 fluorescence, CLN5^KO^ cells had significantly less perinuclear lysosomes compared to wild-type HeLa cells (**Fig. 6I**).

**Figure 6.**
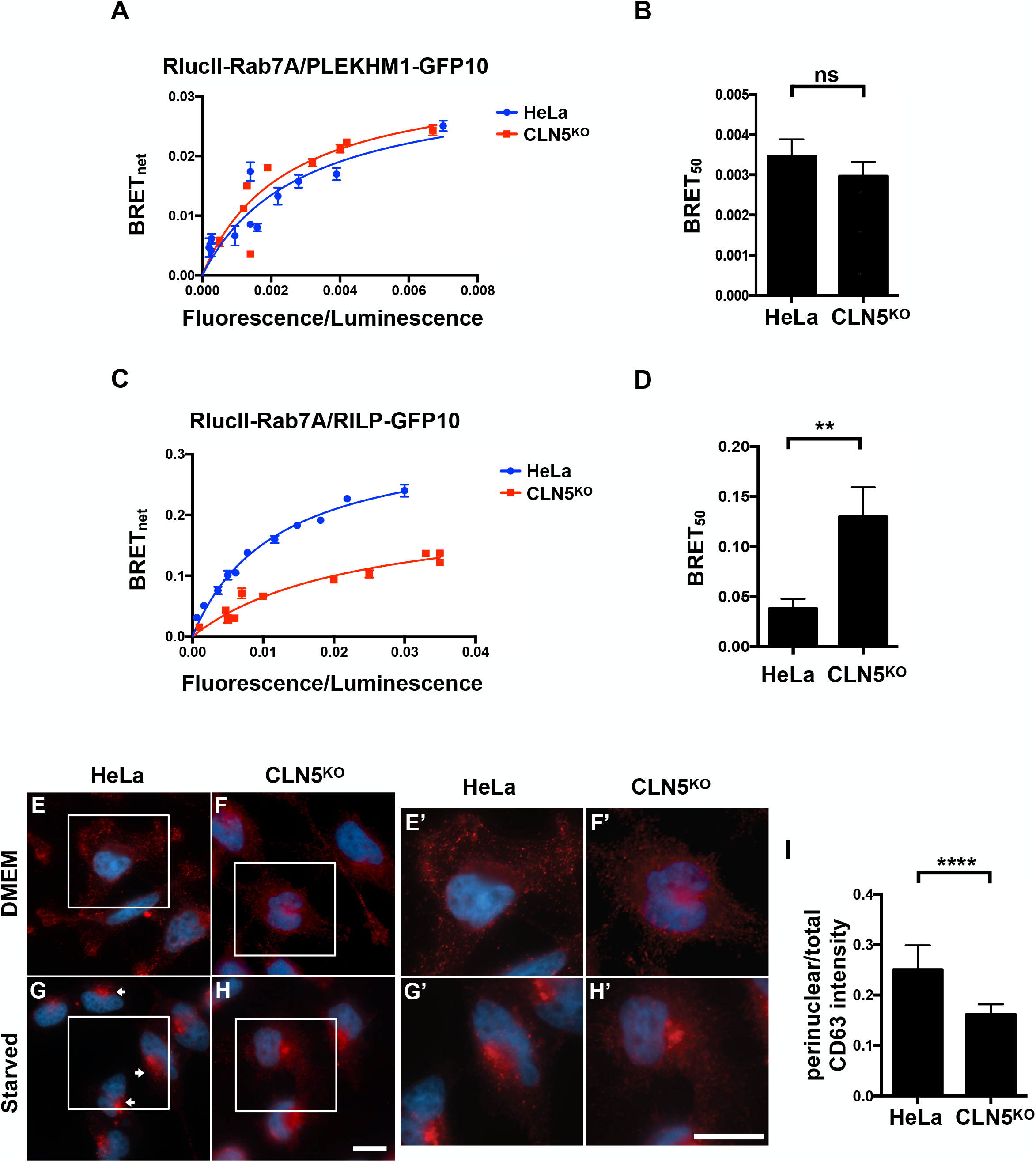
Retrograde transport of lysosomes is deficient in CLN5^KO^ HeLa cells. **(A)** Wild-type and CLN5^KO^ HeLa cells were transfected with a constant amount of RlucII-Rab7A and increasing amounts of PLEKHM1-GFP10 to generate BRET titration curves. BRET signals are plotted as a function of the ratio between the GFP10 fluorescence over RlucII luminescence. BRET_50_ was extrapolated from 3 independent experiments. Data is shown as mean±s.d.; ns, not significant. (**C**) Wild-type and CLN5^KO^ HeLa cells were transfected with a constant amount of RlucII-Rab7A and increasing amounts of RILP-GFP10 to generate BRET titration curves. BRET signals are plotted as a function of the ratio between the GFP10 fluorescence over RlucII luminescence. (**D**) BRET_50_ was extrapolated from 3 independent experiments. Data is shown as mean±s.d.; **P≤0.01, Student’s *t*-test. (**E - H**) Wild-type (**E and G**) and CLN5^KO^ (F and H) HeLa cells were cultured in standard DMEM (**E and F**) or starved (**G and H**) for 3 hours in EBSS. The cells were subsequently immunostained with anti-CD63 antibody. The cells were then fixed with 4% PFA for 12 min, followed by staining with DAPI to visualize the nucleus. Images were taken on a Zeiss Fluorescence microscope using a 63X objective. Scale bar: 10 µm. (**E’ - H’**) Magnified view of white square area shown in E - H. Scale bar µm. (**I**) Fluorescence intensity of CD63 in the perinuclear region and total cellular CD63 fluorescence was determined using ImageJ. Results shown are the ratio of perinuclear divided by total CD63 fluorescence from 30 cells. Data is shown as mean±s.d.; ****P≤0.0001, Student’s *t*-test.

A role for minus-end movement of lysosomes is to enable efficient fusion with autophagosomes, a step required for efficient autophagic degradation [42]. Previous studies have reported autophagic defects in cells depleted of CLN5 by RNAi [43], or CLN5 deficient cells [6, 44]. Therefore, in CLN5^KO^ HeLa cells, we would expect less autophagosome/lysosome fusions and defective autophagic flux. To test fusion ability, we expressed mCherry-LC3 and Lamp1-GFP in wild-type (**Fig. 7A - C**), CLN5^KO^ (**Fig. 7D - F**) and Rab7A^KO^ HeLa cells (**Fig. 7G - I**). LC3 is a marker of autophagosomes, while Lamp1 is a marker of lysosomes. Upon the initiation of autophagy by starvation, LC3-positive autophagosomes fuse with Lamp1-positive lysosomes, forming autolysosomes in order to degrade material [45]. Cells were nutrient starved for 3 hours in EBSS and the co-localization of mCherry-LC3 and Lamp1-GFP was determined. Quantification of the co-localization using Pearson’s coefficient in 30 cells per condition showed a high level of co-localization of mCherry-LC3 and Lamp1-GFP in wild-type HeLa cells, suggesting that autophagosomes were fusing with lysosomes (**Fig. 7J**). Rab7A participates not only in the movement of lysosomes, but also in the fusion process [46]. As such, we would expect significantly less co-localization in Rab7A^KO^ HeLa cells. Compared to wild-type cells, the co-localization between LC3 and Lamp1 was significantly reduced in Rab7A^KO^ HeLa cells (**Fig. 7J**). We also found significantly less co-localization between LC3 and Lamp1 in CLN5^KO^ HeLa cells (**Fig. 7J**), suggesting that the deficient retrograde movement of lysosomes was preventing fusion between autophagosomes and lysosomes.

**Figure 7.**
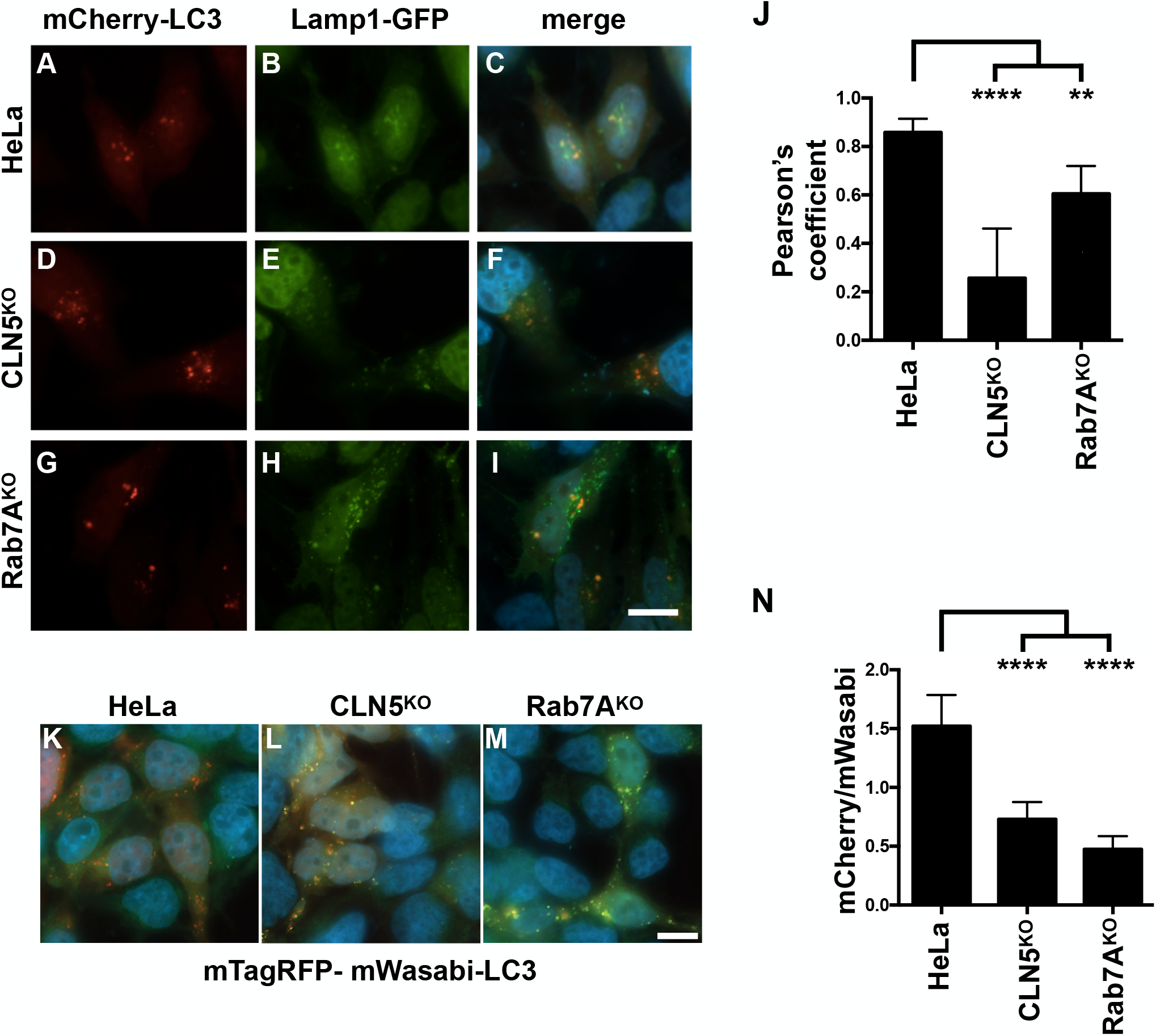
Lack of fusion of autophagosomes to lysosomes in CLN5^KO^ HeLa cells. (**A - I**) mCherry-LC3 and Lamp1-GFP were co-transfected into wild-type (**A - C**), CLN5^KO^ (**D - F**) and Rab7A^KO^ (**G - I**) HeLa cells and imaged on a Zeiss Fluorescence microscope using a 63× objective. Representative images are shown. Scale bar = 10 µm. (**J**) Quantification of the fluorescence from 35 cells to determine the Pearson’s coefficient of mCherry-LC3 and Lamp1- GFP in wild-type, CLN5^KO^ and Rab7A^KO^ HeLa cells. **P≤0.01; ****P≤0.0001; one-way ANOVA followed by Tukey’s post hoc test. (**K - N**) mTagRFP-mWasabi-LC3 was transfected into wild-type (K), CLN5^KO^ (L) and Rab7A^KO^ (M) HeLa cells. Representative images are shown. Scale bar: 10 µm (N) Quantification of mCherry divided by mWasabi was calculated for 30 cells. ****P≤0.0001; one-way ANOVA followed by Tukey’s post hoc test.

Next, we used a tandem LC3 probe (mTagRFP-mWasabi-LC3) to determine autophagic flux [47]. mWasabi is pH sensitive (more sensitive compared to EGFP), while mTagRFP is not. If cells can proceed with autophagy, the fusion of autophagosomes with lysosomes quenches the mWasabi so red is observed, while late blocks in autophagy (no autophagosome fusion with lysosomes) results in an increase in yellow puncta, with a corresponding decrease in red signal, since mWasabi is not quenched. We expressed mTagRFP-mWasabi-LC3 in wild-type (**Fig. 7K**), CLN5^KO^ (**Fig. 7L**) and Rab7A^KO^ HeLa cells (**Fig. 7M**), and the cells were starved in EBSS for 3 hours. In wild-type cells, autophagosome/lysosome fusion occurs, resulting in a red signal as the mWasabi is quenched (**Fig. 7K and N**). In CLN5^KO^ HeLa cells, fusion is deficient, and therefore mWasabi is not exposed to the acidic pH of the lysosomal lumen and no quenching occurs. Therefore, the red signal is significantly decreased (**Fig. 7L and N**). Since Rab7A is required for efficient fusion, a similar observation was made in Rab7A^KO^ HeLa cells as expected (**Fig. 7M and N**).

## Discussion

CLN5 is a soluble endolysosomal protein whose function remains poorly understood. Mutations in this protein cause a rare neurodegenerative disease that affects children, so understanding the function of CLN5 could lead to new therapeutic targets. Work using the model organism *Dictyostelium discoideum* showed that CLN5 has glycoside hydrolase activity [4], and previous reports found that CLN5 regulates lysosomal pH [5] and autophagy [43, 44]. More recent work has implicated CLN5 in mitochondrial function [6]. Using siRNA (CLN5^KD^), we previously demonstrated that CLN5 regulates the stability of the lysosomal sorting receptor sortilin by enabling its endosome-to-TGN trafficking [7]. However, using this KD system, we never tested the effects of disease-causing mutations on this trafficking pathway, or whether CLN5 modulated other Rab7A functions. Furthermore, it is not known how a soluble endolysosomal protein can regulate protein-protein interactions in the cytosol. In this current work, we generated a CLN5 knockout (CLN5^KO^) HeLa cell line that can be rescued with wild-type or mutant CLN5. Using this tool, we have confirmed our previous work and expanded our understanding of the function of this protein.

In our CLN5^KD^ HeLa cells, we observed less membrane bound Rab7A and retromer [7]. We found less membrane bound retromer in our CLN5^KO^ HeLa cells, which was rescued by expressing wild-type CLN5, but not the disease-causing mutant (CLN5^Y392X^). Although retromer recruitment was deficient in CLN5^KO^ HeLa cells, Rab7A was still membrane bound. This raised two points. Why the discrepancy between the KD and KO models, and how could Rab7A be membrane bound but not recruit retromer? We hypothesize that chronic loss of CLN5 in the KO model results in the cell overcoming the effect, while acute depletion of CLN5 by siRNA does not allow time for the cell to compensate. We have previously shown that Rab7A palmitoylation is required for the efficient recruitment of retromer to endosomal membranes, but is not required for Rab7A membrane localization [22]. We found that Rab7A palmitoylation is significantly decreased in CLN5^KO^ HeLa cells, providing an explanation for the decrease in the Rab7A/ retromer interaction as measured in live cells by BRET, and the subsequent lack of retromer recruitment. Palmitoylation of Rab7A is not required to modulate the Rab7A/retromer interaction *per se*, but serves to localize Rab7A to specific endosomal membrane domains to favour the interaction [22]. When membranes are disrupted such as in co-immunoprecipitation, non- palmitoylatable Rab7A can interact with retromer [22]. As such, it is not surprising that Rab7A and retromer interacted by co-immunoprecipitation in CLN5^KO^ HeLa cells.

The recruitment of retromer to endosomal membranes enables the endosome-to-TGN retrieval of the lysosomal sorting receptor sortilin [8]. The sortilin/retromer interaction was perturbed in CLN5^KO^ HeLa cells compared to wild-type HeLa cells. While the expression of wild-type CLN5 in CLN5^KO^ HeLa cells rescued the sortilin/retromer interaction, the expression CLN5^Y392X^ did not. This would suggest that the residues at the C-terminal end of CLN5 are required for the sortilin/retromer interaction, however how those residues modulate this interaction is yet to be determined. The decrease in the sortilin/retromer interaction observed in the CLN5^KO^ HeLa cells should result in the lysosomal degradation of the lysosomal sorting receptor. As in the CLN5^KD^ HeLa cells [7], sortilin is degraded in CLN5^KO^ HeLa cells. Expression of wild-type CLN5 rescued this degradation, while expressing the disease-causing mutant CLN5^Y392X^ did not rescue. Taken together, these observations point to a crucial role of CLN5 in modulating Rab7A-dependent retromer recruitment and function. Indeed by modulating the palmitoylation level of the small GTPase, CLN5 establishes the conditions for an optimal Rab7/ retromer interaction, hence efficient retromer/sortilin binding and recycling to the TGN, which in turns translates in proper lysosomal function.

In our recently published paper, we demonstrated that CLN3 functions as a scaffold to ensure efficient protein-protein interactions. In that study, we demonstrated decreased Rab7A/ retromer and retromer/sortilin interactions in CLN3^KO^ HeLa cells, which consequently lead to the degradation of sortilin (Yasa et al., 2020). We found similar results in our CLN5^KO^ HeLa cells. Furthermore, we found that CLN5 is required for the CLN3/retromer and CLN3/sortilin interactions. Although wild-type CLN5 rescued the CLN3/sortilin interaction in CLN5^KO^ HeLa cells, the expression of the disease-causing mutant did not. Based on this data, we propose a model where CLN3 and CLN5 function as an endosomal complex required for the efficient endosome-to-TGN trafficking of sortilin. This inefficient recycling results in defective lysosomal function, which is a hallmark of CLN5 disease. Our results suggest a potential pathogenic mechanism for CLN5 disease.

We next asked if other Rab7A mediated pathways are affected in CLN5^KO^ HeLa cells. We found significant delays in the degradation of both EGFR and EGF. This delayed degradation could be due to less efficient lysosomal function, or due to decreased fusion of endocytic cargo with lysosomes, a mechanism also modulated by Rab7A. The Rab7A effector RILP is implicated in the degradation of internalized cargo such as EGFR by mediating the movement of vesicles (Wijdeven et al., 2016), while the Rab7A effector PLEKHM1 is a tether required for fusion in both endocytic degradation and autophagy [12]. In CLN5^KO^ HeLa cells, we found no changes to the Rab7A/PLEKMH1 interaction, but we found a decreased Rab7A/RILP interaction. This resulted in deficient movement of CD63 positive lysosomes toward the perinuclear region, an important step in the autophagic process. Indeed, we observed less co- localization of LC3II containing autophagosomes with Lamp1 positive lysosomes upon starvation in CLN5^KO^ HeLa cells, and less autophagic flux as demonstrated using a tandem LC3 probe (mTagRFP-mWasabi-LC3). While decreased lysosomal function plays a role in the pathogenic mechanism of CLN5 disease, defective endocytic degradation and autophagy are most likely due to defective lysosomal movement and thus decreased fusion events.

In conclusion, in this study we demonstrate a role for CLN5, together with CLN3, acting as a complex to regulate endosome-to-TGN trafficking. While CLN3 behaves like an interaction platform, CLN5 ensures those CLN3 interactions, which are vital for lysosomal functioning are maintained and regulated. These findings enlighten the molecular mechanisms behind NCL pathogenesis caused by defected CLN5 and CLN3 proteins.

### Statistics

Statistical analysis was performed using GraphPad Prism Version 7. The statistical tests used are described in the corresponding figure legends.

## Data Availability

All of the primary data that is presented in this study can be requested in electronic form by contacting Stephane Lefrancois. All reagents generated by our group are available upon request to Stephane Lefrancois.

## CRediT Contribution

Stephane Lefrancois: Conceptualization, Supervision, Funding acquisition, Formal Analysis, Writing — original draft, Project administration, Writing — review and editing. Seda Yasa: Conceptualization, Investigation, Methodology, Formal Analysis, Writing — original draft, Writing — review and editing. Graziana Modica: Conceptualization, Investigation, Formal Analysis, Writing — review and editing. Etienne Sauvageau: Investigation, Methodology, Formal Analysis, Writing — review and editing.

## Acknowledgements

We would like to thank Michel Bouvier (IRIC and University de Montreal), Regis Grailhe, (Pasteur Institute Korea), Paul Odgren (University of Massachusetts Medical School, Addgene #73592), Makoto Kanzaki (Tohoku University), Takeshi Nakamura (Tokyo University of Science), Peter J. McCormick (Queen Mary University), Juan S. Bonifacino (NICHD, NIH), Jian Lin (Peking University) for providing plasmids. We would like to thank Guido Hermey (Institute for Molecular and Cellular Cognition, Center for Molecular Neurobiology Hamburg, University Medical Center Hamburg-Eppendorf) for providing the parental HeLa cells.

