## Supplementary material for "CLN5 and CLN3 function as a complex to regulate endolysosome function": Supp Figures

### Yasa et al., 2021 Figure S1

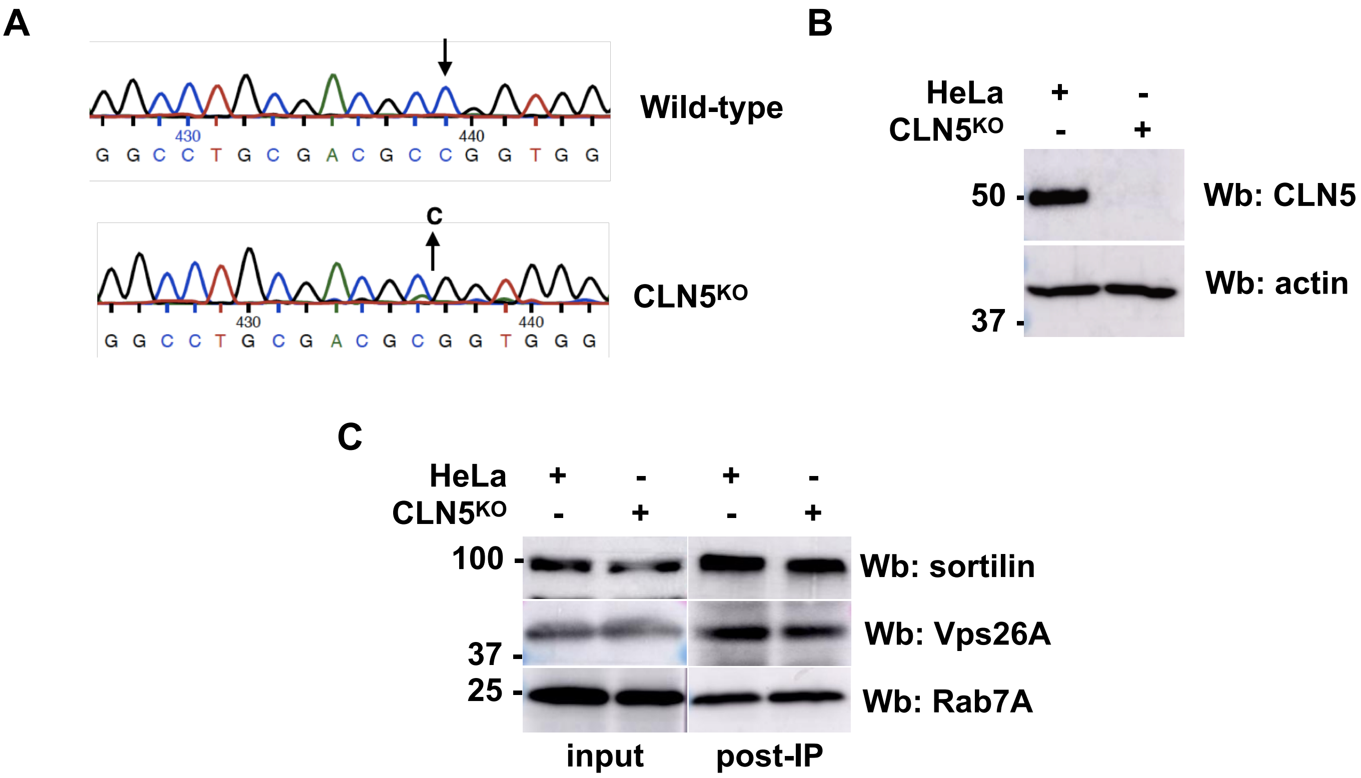


(**A**) DNA sequencing of genomic DNA of wild type and CLN5 knockout (CLN5$^{KO}$) HeLa cells demonstrates the deletion of 1 base pairs in the CLN5 gene. The deletion causes a frameshift, changes the encoded amino acid sequence leading to a premature stop codon after amino acid 47. Arrow indicates the position of the mutation. (**B**) Whole cell lysate from wild-type and CLN5$^{KO}$ HeLa cells was run on an SDS-PAGE and Western blotting (Wb) was performed using anti-CLN5 and anti-actin antibodies. (**C**) Retromer (Vps26A) was immunoprecipitated with anti-Vps26A antibody from wild-type and CLN5$^{KO}$ HeLa cells. Samples were run on a 12% SDS-PAGE and Western blotting (Wb) was performed with anti-sortilin, anti-Vps26A and anti-Rab7A antibodies.

### Yasa et al., 2021 Figure S2

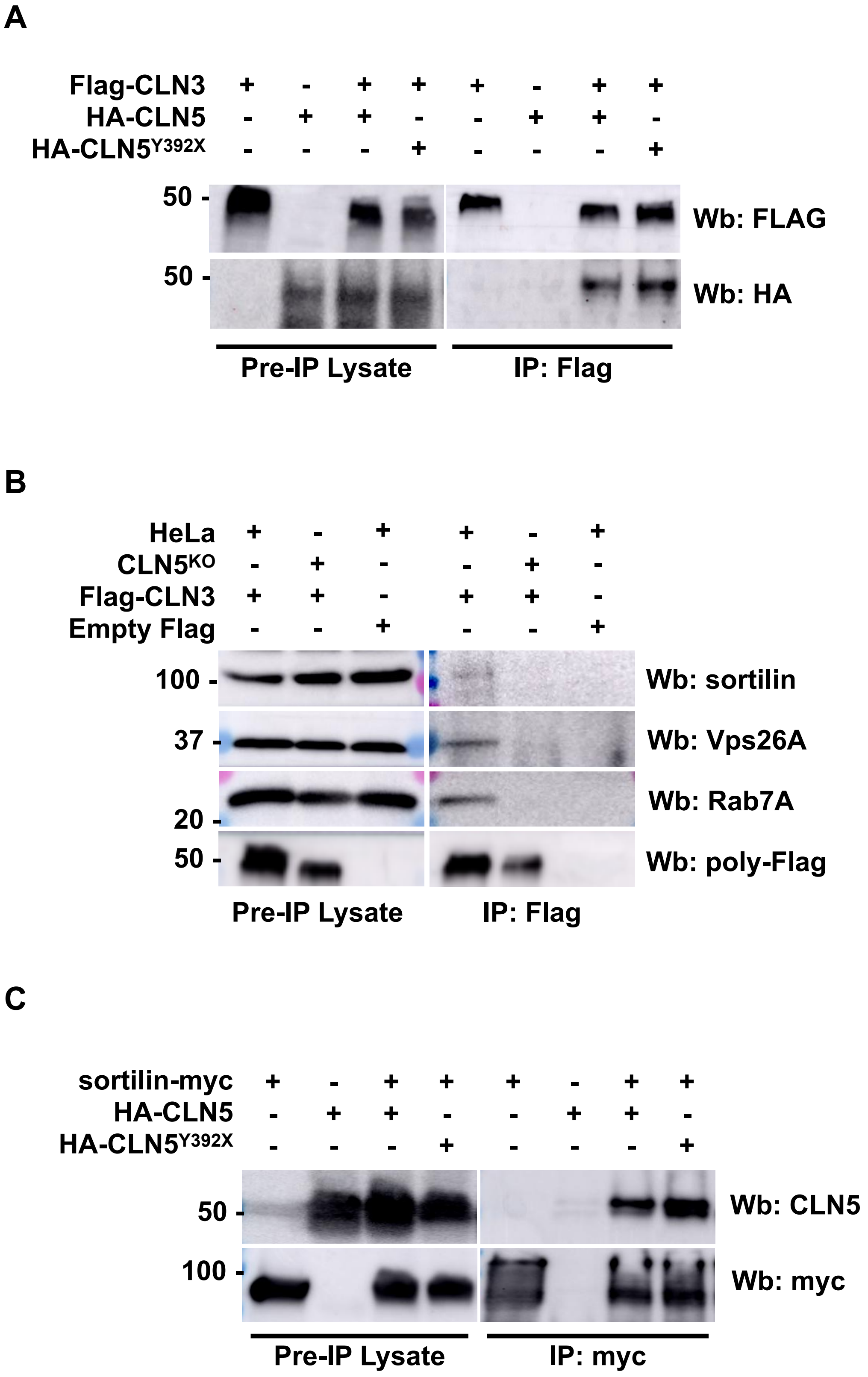

### Yasa et al., 2021 Figure S2

**Figure S2. CLN5 mutations do not affect its interactions**
(**A**) HeLa cells were co-transfected with Flag-CLN3 and wild-type HA-CLN5 or HA-CLN5$^{Y392X}$ as indicated. Flag-CLN3 was immunoprecipitated with polyclonal anti-Flag antibody. Samples were the run on SDS-PAGE and Western blotting (Wb) was performed using monoclonal anti-Flag and anti-HA antibodies. (**B**) Wild-Type and CLN5$^{KO}$ HeLa cells were transfected with Flag-CLN3 or empty plasmid as indicated. Flag-CLN3 was immunoprecipitated with monoclonal anti-Flag antibody. Samples were then run on SDS-PAGE and Western blotting (Wb) was performed using polyclonal anti-Flag, anti-Vps26A, anti-sortilin and anti-Rab7A antibodies. (**C**) CLN5$^{KO}$ HeLa cells were co-transfected with sortilin-myc and wild-type HA-CLN5 or HA-CLN5$^{Y392X}$ as indicated. Sortilin-myc was immunoprecipitated with monoclonal anti-myc antibody. Samples were the run on SDS-PAGE and Western blotting (Wb) was performed using monoclonal anti-myc and anti-CLN5 antibodies.
